# Selective single molecule sequencing and assembly of a human Y chromosome of African origin

**DOI:** 10.1101/342667

**Authors:** Lukas F.K. Kuderna, Esther Lizano, Eva Julià, Jessica Gomez-Garrido, Aitor Serres-Armero, Martin Kuhlwilm, Regina Antoni Alandes, Marina Alvarez-Estape, Tyler Alioto, Marta Gut, Ivo Gut, Mikkel Heide Schierup, Oscar Fornas, Tomas Marques-Bonet

**Affiliations:** Institut de Biologia Evolutiva, (CSIC-Universitat Pompeu Fabra), PRBB, Doctor Aiguader 88, Barcelona, Catalonia 08003, Spain; Institut Hospital del Mar d'Investigacions Mèdiques (IMIM), Carrer del Doctor Aiguader 88, PRBB Building, Barcelona 08003, Spain; Centre for Genomic Regulation (CRG), The Barcelona Institute for Science and Technology, Carrer del Doctor Aiguader 88, PRBB Building, Barcelona 08003, Spain; CNAG-CRG, Centre for Genomic Regulation (CRG), The Barcelona Institute of Science and Technology, Baldiri Reixac 4, Barcelona 08028, Spain; Universitat Pompeu Fabra (UPF), Barcelona, Spain; Bioinformatics Research Center, Aarhus University, C.F. Moellers Alle 8, Aarhus C, Denmark; Department of Bioscience, Aarhus University, Ny Munkegade 116, Aarhus C, Denmark; Institució Catalana de Recerca i Estudis Avançats (ICREA), Passeig Lluís Companys 23, Barcelona, Catalonia 08010, Spain

## Abstract

Mammalian Y chromosomes are often neglected from genomic analysis. Due to their inherent assembly difficulties, high repeat content, and large ampliconic regions^1^, only a handful of species have their Y chromosome properly characterized. To date, just a single human reference quality Y chromosome, of European ancestry, is available due to a lack of accessible methodology^2–5^. To facilitate the assembly of such complicated genomic territory, we developed a novel strategy to sequence native, unamplified flow sorted DNA on a MinION nanopore sequencing device. Our approach yields a highly continuous and complete assembly of the first human Y chromosome of African origin. It constitutes a significant improvement over comparable previous methods, increasing continuity by more than 800%^6^, thus allowing a chromosome scale analysis of human Y chromosomes. Sequencing native DNA also allows to take advantage of the nanopore signal data to detect epigenetic modifications in situ^7^. This approach is in theory generalizable to any species simplifying the assembly of extremely large and repetitive genomes.

---

Recombinational arrest in the common ancestor of the X and Y chromosomes led to the degeneration and accumulation of large amounts of repetitive DNA on the Y chromosome along its evolutionary trajectory^8^. Furthermore, many sequencing efforts have traditionally chosen female samples, as the hemizygous nature of the sex chromosomes leads to half the effective sequencing coverage on both of them in a male, resulting in inferior genome assemblies^9^. Together, these causes have led to an underrepresentation of Y chromosomes in genomic studies and proper characterization of the Y chromosome in only a handful of mammalian species through a time- and labor-intensive clone by clone approach^3–5,10^. One strategy to reduce the complexity of the assembly problem for the Y chromosome, is to isolate it by flow cytometry, thus dramatically reducing the potential amount of overlaps of repetitive regions in the context of the whole genome^1^. Notwithstanding, previous efforts which sought to do this have faced some drawbacks, as the material has been heavily amplified post sorting to increase yield^6^. Whole genome amplification (WGA) introduces biases that are detrimental to genome assembly, such as unequal sequence coverage and chimera formation, as well as limited fragment length^11^. Moreover, these methods lead to the loss of epigenetic modifications that can now be directly determined from the signal data from nanopore sequencers^7^. Additionally, previous efforts to assemble the Y chromosome purely from flow sorted material did so using the gorilla^6^, a species with a previously uncharacterized Y chromosome, meaning that potential biases in the assembly cannot be detected without a gold standard reference to compare to, such as human. Integrating single molecule sequencing has been shown to produce far superior whole genome shotgun assemblies than sequencing by synthesis platforms^12–15^. Furthermore, the MinION sequencing platform from Oxford Nanopore Technologies has recently been used to create the most contiguous human whole genome shotgun assembly to date^16^ and to resolve the structure of the human Y chromosome centromere^17^. To take advantage of these benefits, we developed a protocol to sequence native, unamplified flow-sorted DNA on the MinION sequencing device. We sorted approximately 9,000,000 individual Y chromosomes from a lymphoblastoid cell line (HG02982) from the 1000 Genomes Project, whose haplogroup (A0) represents one of the deepest known splits in humans^18^ (see figure 1a). Given the large volume in which the chromosomes were sorted, and potential issues with residual dyes that are necessary for the sorting process, we devised a purification protocol to bring the DNA into conditions suitable for sequencing. We ran four Oxford Nanopore MinION flowcells to generate 305,528 reads summing to a total of 2,329,124,808 bases of data. The yields per flowcell varied considerably between from 897.6 Mb to 163.8 Mb (see figure 1c). Sequencing yields were on the lower end of the reported spectrum, but read N50 surpassed most of them^16^ (see figure 1d). Additionally, for the same flow sorted material we ran an Illumina MiSeq lane for 2x300 cycles but including 4 rounds of PCR amplification. To check the enrichment specificity, we aligned the reads to the human reference genome (GRCh38) and calculated the normalized coverage on each chromosome. Taking into account the size of the Y chromosome and its haploid nature, we find it to be over 110-fold enriched compared to a random sampling from the human genome (see figure 1b and supplementary material).

**Figure 1:**
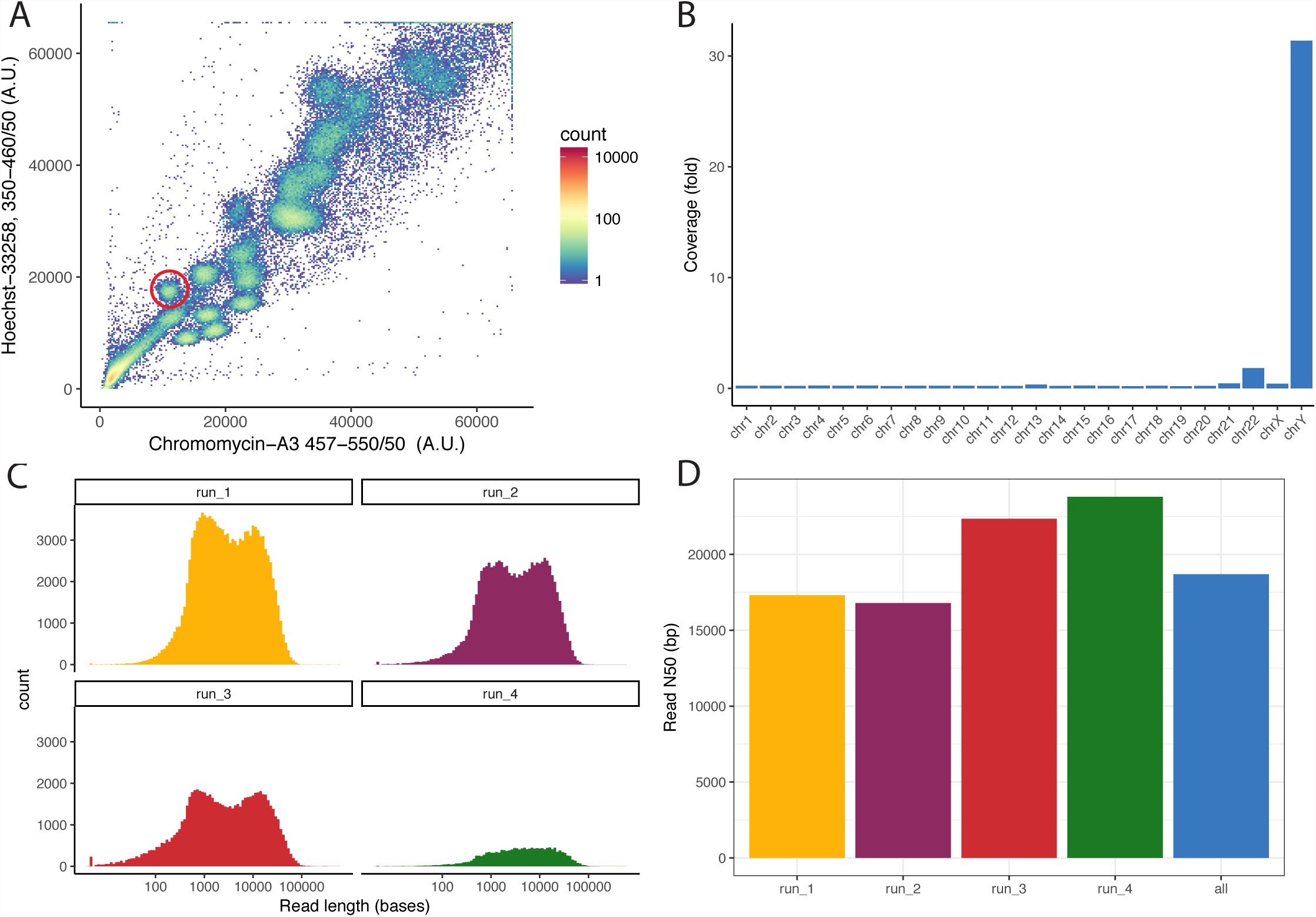
**A** Flow-karyogram of a human genome. The different clusters correspond to different chromosomes. The red circle delimits the cluster corresponding to the Y chromosome used for this project. **B**: Enrichment specificity of the sequencing data. Sequences on the Y chromosome are ~110-fold enriched compared to WGS sequencing. Chromosome 22 partially co-sorts with Y. All other chromosomes are depleted. **C**: Read-length distribution of the four runs. **D**: N50 values for all four runs and the combined dataset.

We used the Nanopore data to construct a complete *de novo* assembly using Canu^19^. We performed a self-correction by aligning the reads used for assembly and called consensus using Nanopolish^7^, correcting a total of 127,809 positions. Finally, the Illumina library served to polish residual errors within the assembly using pilon^20^. By this means, we corrected a further 101,723 single nucleotide positions and introduced 105,640 small insertions and 6,983 small deletions. The final assembly is comprised of 35 contigs, with an N50 of 1.46 Mb amounting to 21.5 Mb, in contrast to a contig N50 of 6.91 Mb of the GRCh38 Y chromosome assembly. Compared to the gorilla Y chromosome assembly with a contig N50 of 17.95 kb^6^, our assembly is two orders of magnitude more contiguous (see figure 2b).

The Y chromosome is comprised of a set of discrete sequence classes^10^. To check the completeness of our assembly we assessed how well each of them is represented. After retaining only single best placements, we were able to align 21.2 Mb, or 98.9% of its total length, with 99.9% of identical bases on average (see figure 2a). We recovered the full-length (>99% of the annotated length in GRCh38) reconstructions of both the X-transposed and the X-degenerate regions. While the X-degenerate region can be considered a single copy region due its distant common ancestry with the X chromosome, the X-transposed region emerged only after the split between humans and chimpanzees^21^. The largest sequence class on the Y chromosome is comprised of ampliconic regions, which amount to around 30% of the euchromatic portion and sum to 9.93 Mb. These regions contain eight massive, segmentally duplicated palindromes, all of which share more than 99.9% identity between their two copies, with the largest one spanning over 2.90 Mb. We find this region to be the most challenging to reconstruct, with fragmented and collapsed sequences, but are nevertheless able to recover 6.44 Mb, or 65.6% of its length in GRCh38. Surprisingly, we recover only 80.2% of the pseudo-autosomal regions (PARs). We observed a rather steep drop-off in coverage coinciding with the PAR-1 boundary on GRCh38. As we are sequencing native, unamplified DNA, the genomic coverage is directly proportional to the number of copies of the underlying sequenced region^22^. We compared the mapped coverage of our raw data on GRCh38 and find that PAR-1 exhibits only around 72% of the average coverage of the whole chromosome (19.8-fold versus 27.3-fold). Given the sharp coincidence with the PAR-1 boundary, we suspect this to not be a technical artifact of the library preparation (see supplementary material). Finally, of the remaining sequence classes we are able to recover around 28.8% of the resolved heterochromatic regions, and 98.9% of the remaining unclassified sequences (referred to as “other”; see Table 1).

**Figure 2:**
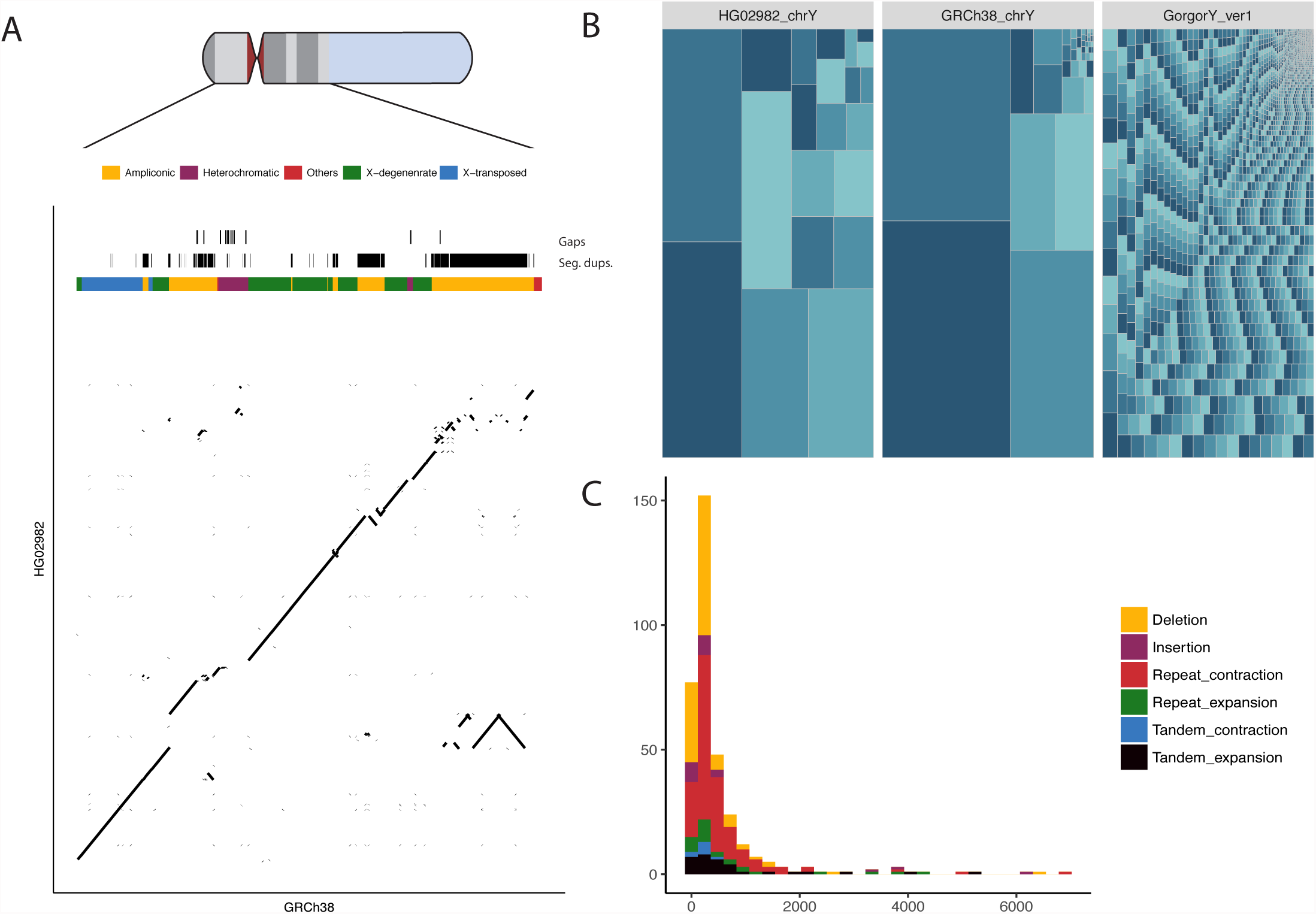
**A** Dot-plot comparing GRCh38 to HG02982. The reconstruction is highly continuous along most sequences classes, with ampliconic regions showing a higher degree of fragmentation. **B**: Treemap comparing the contiguity of HG02982 chrY to GRCh38 chrY and the gorilla Y chromosome by Tomaszkiewicz et al. The size of each rectangle corresponds to the size of a contig within each of the assemblies. **C**: Histogram of size distribution of the structural variants called against GRCh3 8.

**Table 1:** Summary of sequence class coverage of HG02982 versus GRCh38, as well as the contigs from NA23385 identified as derived from the Y chromosome.

| Sequence Class | Aligned Length<br>HG02982<br>(bp) | Identity of alignments<br>HG02982<br>(%) | Proportion recovered in<br>HG02982 from<br>GRCh38 (%) | Aligned Length<br>NA24385 WGS<br>(bp) | Proportion recovered in<br>NA24385 WGS<br>from GRCh38<br>(%) | Length of seq. class<br>in GRCh38 w/o<br>gaps (%) |
| --- | --- | --- | --- | --- | --- | --- |
| **Ampliconic** | 6,437,254 | 99.93 | 65.64 | 5,242,461 | 53.46 | 9,807,089 |
| **Heterochromatic** | 477,140 | 99.69 | 28.80 | 171,045 | 10.32 | 1,656,797 |
| **Others** | 75,675 | 99.04 | 98.86 | 63,973 | 83.57 | 76,547 |
| **Pseudo-autosomal** | 2,283,873 | 99.71 | 80.28 | 117,626 | 4.13 | 2,844,939 |
| **X-degenerate** | 8,573,300 | 99.94 | 99.36 | 8,238,733 | 95.48 | 8,628,904 |
| **X-transposed** | 3,368,361 | 99.94 | 99.05 | 1,474,610 | 43.36 | 3,400,750 |

To contrast our approach to a long-read whole genome shotgun (WGS) assembly, we assembled the publicly available PacBio dataset from the Ashkenazim son from the Genome in a Bottle Consortium^23^, which has a sequencing depth comparable to ours on the sex chromosomes (~30X). We identified 193 contigs mapping to the Y chromosome, with an N50 of 213 kb, covering 15.3 Mb, or around 28% less than by our approach. The WGS fails to assemble roughly a 56.6% of the X-transposed region and 47% of the ampliconic regions (see Table 1). We hence show our approach to yield significantly superior Y-chromosomal reconstruction than from long-read WGS assemblies in terms of content and contiguity.

We performed a comparative annotation to check the completeness of our assembly at the gene level. To this end, we projected all Gencode (v. 27, GRCh38) annotations on the Y chromosome onto our assembly and annotated them there. Due to its peculiar evolutionary trajectory, the gene-space on the Y chromosome is degenerate, and any remaining genes can generally be classified into two categories: On one hand there are single copy genes, which are broadly expressed beyond the testis. On the other, there are multi-copy genes within the ampliconic regions, that are mainly involved in spermatogenesis^24^. We recover the complete gene set of the genes in the MSY region and are therefore able to annotate all single copy genes. Furthermore, we are able to retrieve at least one member of all multi-copy gene families. For 4 out of 9 of these gene families, we are additionally able to resolve further copies within our assembly. Surprisingly, we find four genes (ASMTL, IL3R, P2RY, SLC25) from a comparatively short syntenic block of around 200 kb in the PAR-1 region to be at least partially missing from our assembly. However, mapping the raw data onto GRCh38 shows that this is an artifact, and the deletion is due to the aforementioned assembly difficulties within the PAR-1 region.

We produced a stringent call set of structural variants (SVs) derived from alignments to GRCh38 using Assemblytics^25^. We detect 347 SVs at least 50 bp in size (931 variants at least 10 bp in size) of which 82 are at least 500 bp long (see Figure 2C). The cumulative length of these variants sums to 184 kb. We observe a 4.8-fold excess number of deletions versus number of insertions, amounting to a 2-fold excess of bases in deletions versus bases in insertions. While a deletion bias for nanopore based assemblies had previously been reported^16^, we find the strength of this bias to be decreasing in our analysis, probably reflecting improvements in base-calling accuracy. To check the presence of large-scale copy number variation in multi-copy genes, we additionally determined the chromosome-wide copy number based on a read-depth approach. We find extensive genic copy number variation, with expansions in 5 of the 9 multicopy genes, when compared to the reference individual. Among these, we find expansions in RBMY, PRY, BPY2 and DAZ, all members of the AZFc-region locus with implications for male fertility. Lastly, to assess concordance with previous studies, we compared our SV calls to those generated by the 1000 Genomes Project, which contains the same cell line used for this study^26^. We manually validate all structural variants called in HG02982 in the 1000 Genomes Project in our assembly (see supplementary material).

Finally, we called the methylation status of 5-methylcytosin (5-mc) at CpG positions from the signal data using Nanopolish^7^. We checked the concordance of these calls against whole genome bisulfite sequencing calls from similar primary cell types of the ENCODE project (B-cell, T-cell and natural killer cell), as well as another male cell line derived from skin fibroblast, and find all methylation calls on the Y chromosome to be well correlated with these data (Pearson’s r 0.502-0.583, see supplementary material), and note the previously observed lower methylation levels in LCL in our cell line compared to primary tissues^27^.

Here, we report the first successful sequencing and assembly of native, flow-sorted DNA on an Oxford Nanopore sequencing device, without previous amplification. We apply our methodology to assemble the first human Y chromosome of African origin to benchmark our approach. This is arguably the most challenging human chromosome to assemble due to its high repeat and segmental duplication content, and hence a good test-case to explore the possibilities and limitations of this approach. With the exception of BAC-based assemblies, we are able to reconstruct the Y chromosome to unprecedented contiguity and completeness in terms of sequence class representation. We show that we not only outperform previous efforts that sought to achieve a similar goal^6^, but also accomplish a much better reconstruction than the Y chromosomal sequences derived from a long-read whole genome shotgun assembly. Additionally, our method is orders of magnitude cheaper than reconstructions from WGS data too, especially considering that twice the desired Y chromosomal target coverage is needed on the autosomes. Given the current developments in sequencing throughput, a single MinION flowcell should now be sufficient to assemble a whole human Y chromosome. Furthermore, it is becoming clear that the upper read length boundary is only delimited by the integrity of the DNA, suggesting the possibility that complete Y chromosome assemblies, including full resolution of amplicons, might be possible in the near future. It also is worth noting that our efforts to sequence the same input material on Pacific Biosciences Sequel platform have been fruitless, presumably due to interference of residual dyes with the sequencers optical detection system. The method described here offers the opportunity to take advantage of the benefits of long-range data together with local complexity reduction. Immediate applications are either very complex chromosomes, such as the human Y, or extremely large genomes with a very high degree of common repeats which have long challenged traditional whole genome shotgun approaches, such as wheat, the loblolly pine, or the axolotl^28–31^.

## Acknowledgements

This study was supported by the Spanish Ministry of Economy and Competitiveness with Proyectos de I+D “Excelencia” y Proyectos de I+D+I “Retos Investigación” BFU2014-55090-P awarded to T.M-B and O.F, Centro de Excelencia Severo Ochoa 2013-2017 and Centro de Excelencia Maria de Maeztu 2016-2019. We acknowledge the support from the CERCA Programme of the Generalitat de Catalunya, institutional support from the Spanish Ministry of Economy, Industry and Competitiveness (MEIC) through the Instituto de Salud Carlos III, from the Generalitat de Catalunya through the Departament de Salut and Departament d’Empresa i Coneixement, and co-financing by the Spanish Ministry of Economy, Industry and Competitiveness (MEIC) with funds from the European Regional Development Fund (ERDF) corresponding to the 2014-2020 Smart Growth Operating Program. L.F.K.K. is supported by an FPI fellowship associated with BFU2014-55090-P (MINECO/FEDER, UE). M.K. is supported by a Deutsche Forschungsgemeinschaft (DFG) fellowship (KU 3467/1-1). TMB is supported by BFU2017-86471-P (MINECO/FEDER, UE), U01 MH106874 grant, Howard Hughes International Early Career, Obra Social “La Caixa” and Secretaria d’Universitats i Recerca del Departament d’Economia i Coneixement de la Generalitat de Catalunya.

## Materials and Methods

### Chromosome preparation for flow karyotyping

Mitotic chromosomes suspension were prepared as previously described^32^ with some modifications. Briefly, lymphoblastoid cell lines were cultured in RPMI 1640 medium supplemented with 2mM L-glutamine (Invitrogen, ref. 21875-034), 15% fetal bovine serum and antibiotics (Penicillin and Streptomycin (Invitrogen, ref. 15140-122)) at initial concentration no less than 150,000 viable cells/ml. Near confluence, cells were subcultured to 50%. After 24h, the cells were blocked in mitosis by adding Colcemid to the culture (10 μg/ml demecolcine solution (Gibco, ref. 15210-040)) to a final concentration of 0.1 μg/ml and incubated for an additional 6-7 h. To swell and stabilize mitotic cells, they were centrifuged 5 min at 300×g at room temperature. The pellet was slowly resuspended in 10 mL hypotonic solution (Hypotonic solution: 75 mM KCl, 10 mM MgSO4, 0.2 mM spermine, 0.5 mM spermidine. pH 8.0), incubated for 10 min at room temperature. After the incubation in the hypotonic solution, the swollen cells were centrifuged at 300×g for 5 min. The cell pellet was resuspended in 1.5 mL of ice-cold polyamine isolation buffer (PAB: 15 mM Tris, 2 mM EDTA, 0.5 mM EGTA, 80 mM KCl, 3 mM dithiothreitol, 0.25% Triton X-100, 0.2 mM spermine, 0.5 mM spermidine. pH 8.0) for 20 min to release the chromosomes. To ensure the integrity of the chromosomes, their morphology was checked before staining them. To this end, the pellet was vigorously vortexed for 30 s to liberate the chromosomes from the mitotic cells. The suspension was filtered through a 35 μm mesh filter and stored at 4°C until its sorting.

Finally, chromosomes were stained with chromomycin-A3 (Sigma, ref. C2659) and Hoechst 33258 (Invitrogen, ref. H3569) at a final concentration of 40 µg/ml and 5 µg/ml respectively in presence of divalent cations (10 mM MgSO4 (Sigma, ref. 60142)). Staining was performed for at least 8 h at 4°C, to allow the dyes to equilibrate. Before the sample analysis on a cell sorter, potassium citrate was added to a final concentration of 10 mM (Sigma, ref. 89306) to enhance peak resolution in the flow karyotype.

### Chromosome sorting

Flow Karyotyping for chromosome sorting was performed on BD Influx cell sorter (Becton Dickinson, San Jose, CA), a jet-in-air cell sorter that was selected for its relatively easy manual daily fine-tuning and high-resolution capabilities. Of the five available lasers, only the blue (488 nm laser at 200 mW), deep-blue (457 nm laser at 300 mW) and ultraviolet (355 nm laser at 100 mW) ones were used for flow karyotyping. The setup and performance were optimized using standard 8-peaks Rainbow beads (Sphero™ Rainbow Calibration Particles 3.0 – 3.4µm, BD Biosciences, ref. 559123), 1-peak UV beads for UV laser alignment (Alignflow™ Flow Cytometry Alignment 2.7µm, Molecular Probes, ref. A16502) and 1-peak 457nm for deep-blue laser alignment (Fluoresbrite™ Plain YG Microspheres 1.0µm, Polysciences, Inc ref. 17154) were respectively used for 488-blue, 355-UV and 457-DeepBlue optimal laser alignment and instrument fine-tuning to obtain the highest resolution of chromosome detection and sorting.

The threshold for chromosome sorting was set triggering in chromomycin-A3 fluorescence on 457nm laser as primary excitation line and set at approximately 1800 a.u. Then, chromomycin-A3 fluorescence was used as primary fluorescence reference through a light line of 500 LP filter and collected by a 550/50nm band-pass filter. Hoechst was excited with the UV laser and its fluorescence was collected through a light line of 400 LP filter and by 460/50 BP. All parameters were collected in lineal mode and analyzed with the BD FACS™ Sortware (v. 1.0.0.0.650, Becton Dickinson, San Jose, CA).

We chose a 100 µm nozzle because we found it to have the best piezoelectric-frequency/electronic-noise ratio. The piezoelectric frequency was adjusted at 38.7 KHz. The sample flow rate for chromosome sorting was adjusted at up to 6000 events s^-1^. The gating strategy for chromosome sorting was simple because only a bi-parametrical dot plot Hoechst vs. chromomycin-A3 fluorescence was used (see Figure 1A).

### Purification and concentration of flow-sorted Y chromosomes

For each of the two rounds of purification, the fractions corresponding to approximately 4.5 M Y chromosomes (~500 ng of DNA per aliquot) were divided into 1 ml aliquots with an estimated chromosome count of 400,000, corresponding to a DNA concentration of approximately 0.04 ng/ul. The approximate total volume per round of purification was around 22.5 ml. Each tube containing the flow-sorted DNA was treated overnight with 10 µl of proteinase K (20 mg/ml) at 50 °C. After treatment, the buffer was exchanged, and proteinase K as well as chromomycin-A3 and Hoechst 33258 removed by dialysis against 1 L of TE buffer using a Pur-A-Lyzer™ Maxi Dialysis column with a MWCO of 50 kDa (Sigma-Aldrich). Dialysis was carried out for 48 h exchanging the buffer every 10-16 h. To reduce the volume after buffer exchange, DNA was transferred into 1.5 ml tubes and concentrated by evaporation in a miVac DNA concentrator (Barnstead GeneVac, Ipswich, UK) up to a volume of approximately 5 to 10 µl. A final purification step was performed by pooling the concentrated DNA into two tubes and subjecting it to a SPRI bead purification with a 2X ratio (SPRI beads/sample). DNA was eluted in 9 μl of Low TE buffer and pooled into one tube. Concentrations were determined by absorbance at 260 nm with a NanoDrop 2000 (Thermo Scientific) and by fluorometric assay with the Qubit 2.0 using the Qubit dsDNA HS kit (Invitrogen) (see supplementary material).

### Sequencing the flow sorted chromosomes

The purified DNA was prepared for sequencing following the protocol in the Rapid Sequencing kit SQK-RAD002 (ONT, Oxford, UK). Briefly, approximately 200 ng of purified DNA was tagmented for 1 min at 75°C with the Fragmentation Mix (ONT, Oxford, UK). The Rapid Adapters (ONT, Oxford, UK) were added along with Blunt/TA Ligase Master Mix (NEB, Beverly, MA) and incubated for 30min at RT. The resulting library was combined with RBF (ONT, Oxford, UK) and Library Loading Beads (ONT, Oxford, UK) and loaded onto a primed R9.4 Spot-On Flow cell (FLO-MIN106). Sequencing and initial base-calling was performed with a MinION Mk1B v1.7.10 software package running for 48 h. Estimates for DNA quantification were based on chromosomal counts with corresponding quantification values from Gribble et al, 2009^32^. The uncertainties in quantification with Qubit 2.0 or NanoDrop are presumed to be due to residual intercalating dyes present within the sample, that interfere with the quantification platforms detection systems, with competition of additional intercalants leading to underestimation on the Qubit 2.0, and the additional presence of aromatic groups leading to overestimation on the NanoDrop.

A total estimated amount of 100 ng of Y chromosome was fragmented on a Covaris ultrasonicator with settings targeting fragments of 450 bp. The library was prepared using NEBNext Ultra II DNA Library Prep Kit (New England BioLabs) following the manufacturer’s instructions, including 4 cycles of PCR amplification. Agilent BioAnalyzer High-Sensitivity DNA Kit was used to determine the size distribution and molarity. The library was sequenced on an Illumina MiSeq using the v3 kit and 600 cycles resulting in 300 bp paired end reads.

### Assembly, error-correction and polishing

The Nanopore data was assembled with Canu (v 1.6)^19^ without previous read separation of reads deriving from different chromosomes and assuming a chromosome size of 52 Mb. The following parameters were used:

~~~
canu -p HG02982 -d HG02982_canu genomeSize=52m overlapper=mhap utgReAlign=true -
nanopore-raw raw_data/HG02982/all.joint.fastq
~~~

The 2.3 Gb of input data resulted in 25X of error corrected reads for assembly, assuming a chromosome size of 52 Mb. The data assembled into 35 contigs which where self-corrected using the Nanopore input reads. To this end, we re-performed base-calling from the fast5 files using Albacore (v 2.1, available from the nanopore user community) to be used for variant calling with Nanopolish (v. 0.8.4, https://github.com/jts/nanopolish, Dec. 11, 2017)

~~~
read_fast5_basecaller.py -f FLO-MIN106 -k SQK-RAD002 -i input_folder -s outout_folder
-t 8 -o fastq,fast5 -q 10000000 -n 100000 --disable_pings
~~~

We indexed the reads to be used with Nanopolish:

~~~
nanopolish index -f fast5.fofn reads.joint.fastq
~~~

The reads were mapped onto the raw assembly using bwa mem (v. 0.7.120)^33^ with the additional flag -x ont2d and the mappings merged and sorted with samtools (v. 1.5):

~~~
bwa mem -x ont2d HG02982_canu. uncorrected. fasta reads. joint. fastq | samtools sort -o
reads.joint.mappings.bam -T tmp -
~~~

The mappings were fed to Nanopolish and corrected in chunks of 50 kb using the helper script ‘nanopolish_makerange.py’ included in the Nanopolish package. Variants were called using ‘nanopolish variants –consensus’ with the optional flag ‘--min-candidate-frequency 0.1’.

~~~
nanopolish_makerange.py HG02982_canu. uncorrected. fasta | xargs -i echo nanopolish
variants --consensus selfcorrected.{}.fa -w {} -r reads.joint.fastq -b
reads.joint.mappings.bam -g HG02982_canu.uncorrected.fasta -t 4 --min-candidate-
frequency 0.1 | sh
~~~

By this means, we corrected 127,801 positions in the initial assembly. The self-corrected assembly was further polished with the Illumina library. To this end, we trimmed the Illumina reads to get rid of any adapters in the sequences using trimgalore (v 3.7, https://github.com/FelixKrueger/TrimGalore).

~~~
trim_galore --fastqc --paired --retain_unpaired –gzip pair1.fastq pair2.fastq
~~~

The trimmed reads were mapped with BWA mem (v.0.7.12)^33^ in paired end mode and the mappings converted to a sorted bam files using samtools sort. PCR duplicates were removed with Picardtools (v. 2.8.2, https://broadinstitute.github.io/picard/).

~~~
bwa mem HG02982_canu.selfcorrected.fasta reads.p1.fastq reads.p2.fastq | samtools sort
-o reads.paired.mappings.bam -T tmp -;
java -jar picard.jar MarkDuplicates I=reads.paired.mappings.bam
O=reads.paired.mappings.markdup.bam M=reads.paired.mappings.markdup.bam
~~~

Polishing was performed with Pilon (v 1.22)^20^, resulting in 132,336 residual errors being corrected.

~~~
java -Xmx96G -jar pilon-1.22.jar --threads 12 --genome HG02982_canu.sel
fcorrected.fasta --frags reads.paired.mappings.markdup.bam --output
HG02982_canu.selfcorrected.pileon --outdir pilon_corrections --changes --vcf --tracks
--fix all
~~~

The PacBio data from the Ashkenazim Son (Coriel ID NA24385) produced by the genome in a bottle consortium was also assembled using Canu (v. 1.6) with default assembly parameters and assuming a genome size of 3.2 Gb:

~~~
canu -p NA24385 -d NA24385_canu genomeSize=3.2g -pacbio-raw data/fastq/*fastq.gz
gridOptionsExecutive=‘--mem-per-cpu=16g --cpus-per-task=2’
~~~

### Variant calls

For variant calls, the Illumina data was mapped onto the GRCh38 or the HG02982 assembly respectively, and processed the same way as detailed above. Variants were called using GATKs Haplotype Caller with the following optional flags: ‘--genotyping-mode DISCOVERY --sampleploidy 1’.

~~~
java -jar gatk-package-4.0.0.0-local.jar HaplotypeCaller -R reference.fa -I
mappings.bam --genotyping-mode DISCOVERY -O variants.vcf
~~~

### Repeat annotations

Repeat annotations were performed using RepeatMasker (v. 4.0.7) with rmblastn v. 2.6.0+ as the engine. To be comparable, the annotations for both the HG02982 as well as the GRCh38 assembly were performed the same way. We used the RepBase-20170127 as the repeatmasker database, and Homo sapiens as the query species. Divergence of the repeat annotations to their consensus was calculated using the ‘calcDivergenceFromAlign.pl’ utility included in the RepeatMasker package.

~~~
RepeatMasker -e ncbi -pa 12 -s -species human -no_is -noisy -dir ./outDir -a -gff -u
reference.fa
~~~

### Whole genome alignments

Whole genome alignments to GRCh38 were produced using last (v. 914) with the following parameters as suggested by the developer for highly similar genomes for indexing and alignments:

~~~
lastdb -uNEAR -R01 index reference.fa
lastal -v -j4 index query.fa > mappings.maf
~~~

Single best placements of query sequences were retained using the ‘last-split’ script included in the last alignment package. Alignments were filtered for a maximum mismap probability of 10e-5. The alignments were converted to psl format for further processing.

### Comparison to WGS PacBio data

The PacBio data from the Ashkenazim Son (Coriel ID NA24385) produced by the genome in a bottle consortium was also assembled using Canu (v. 1.6) using default assembly parameters and assuming a genome size of 3.2 Gb:

~~~
canu -p NA24385 -d NA24385_canu genomeSize=3.2g -pacbio-raw data/fastq/*fastq.gz
gridOptionsExecutive=‘--mem-per-cpu=16g --cpus-per-task=2’
~~~

After genome assembly, we performed a whole genome alignment to GRChg38 and retained single best placements as mentioned above. To identify contigs belonging to the Y chromosome, we performed the following filtering steps: For contigs which have local best placements on a chromosome different than the Y, we filtered out those whose proportion of mapped bases is higher on a sequence from the reference assembly different from the Y chromosome. Additionally, we filtered out any alignments with a mismap probability higher than 10e5. By this means, we retained 193 contigs mapping 15,308,468 base pairs on the Y chromosome (see supplementary material)

### Structural variant calls

Structural variants were called with assemblytics^25^. To this end, we produced whole genome alignments using nucmer from the Mummer package (v. 3.22)^34^. The resulting delta file was passed to assemblytics, with the required unique anchor length set to 10000 bp.

~~~
nucmer -maxmatch -l 100 -c 500 GRCh38.chrY.fa HG02982_chrY_v1.fasta -prefix
HG02982_vs_HG38
Assemblytics HG02982_vs_HG38_.delta HG02982_chrY_v1.vs.hg38_10kanchor.50kmax 10000
bin/Assemblytics/
~~~

### Read depth duplication detection

We estimated absolute copy number using the Illumina data as previously described^22^. Briefly, we masked all common repeats as identified by RepeatMasker (see above) and tandem repeat finder. We created non-overlapping 36-mers of the raw reads, that were mapped onto the assembly using GEM (v 2) allowing for a divergence of up to 5%. The read depth was calculated in non-overlapping windows of 1 kb of non-repetitive sequence. After correcting for GC content using mrCanavar (v. 0.51), we normalized by the mean read depth. To assign a copy number to each gene, we calculated the median copy number of all windows intersecting a gene. For the hg38 Y chromosome, a set of custom single-copy regions needed to be provided to the CN caller as calibration. These regions were inferred by subtracting the reference WGAC (whole genome assembly comparison, UCSC track genomic superdups) segmental duplication track from the whole Y chromosome and keeping only stretches of single-copy sequence longer than 2 kb.

### Gene annotation

The annotation of the HG02982 assembly was performed by trying to assign the genes present in the Y chromosome annotation of GRCh38 gencode version 27. For this purpose, we downloaded the gff3, the transcript sequences and the protein sequences that corresponded to the Y chromosome annotation and performed transcript and protein mappings with GMAP (v. 20170317, ^35^) and exonerate (v. 2.2.0, ^36^), respectively. Additionally, a numeric index was assigned to each gene in the HG38 Y chromosome according to the order in the chromosome. Next, we combined all the data (transcript mappings, protein mappings and gene synteny) with an in-house script to locate each gene in our assembly and assign parts of the assembly to their corresponding region in the Y chromosome of GRCh38. After following the strategy mentioned above for all the genes, we took a closer look to the protein-coding genes, by manually checking some of the mappings in order to determine possible errors in the sequence caused by the Nanopore reads that could introduce frameshifts or internal stop codons in the aminoacidic sequence.

### Comparison of methylation status

The methylation status was called using Nanopolish^7^ as suggested by the developers. To this end, we aligned the Nanopore reads to GRCh38 with minimap2^37^ and sorted with samtools (v 1.5). The calls were performed in 200 kb windows.

~~~
minimap2 -a -x map-ont chrY.fa joint_reads.fastq | samtools sort -T tmp -o
joint_reads.mappings.bam
samtools index joint_reads.mappings.bam
nanopolish call-methylation -v --progress -t 8 -r joint_reads.fastq -b
joint_reads.mappings.bam -g chrY.fa -w”chrY:$start-$stop” > methylation_calls.tsv
~~~

Finally, we calculated the methylation frequency and log-likelyhood ratios of methylation at each position:

~~~
calculate_methylation_frequency.py -i methylation_calls.tsv
~~~

We downloaded publicly available methylation calls based on whole genome bisulfite sequencing (WGBS) from the ENCDODE project to compare to. These included calls from similar primary tissue: b-cell, ID: ENCFF774VLD; t-cell, ID: ENCFF355UVU; killer cell, ID: ENCFF435ETE. Additionally, we also compared our calls to another male cell line derived from skin fibroblasts, ID: ENCFF752NXS. The comparisons were performed using the compare_methylation.py script included in the Nanopolish package. We filtered out any position with less than 10 reads in either the WGBS or the Nanopore data. Additionally, any position with a log-likelihood ratio of less than 2.5 in the Nanopore data was also excluded.

## References

1. Tomaszkiewicz, M., Medvedev, P. & Makova, K. D. Y and W Chromosome Assemblies: Approaches and Discoveries. Trends Genet. 33, 266–282 (2017).

2. Skaletsky, H. et al. The male-specific region of the human Y chromosome is a mosic of discrete sequence classes. Nature 423, 825–837 (2003).

3. Hughes, J. F. et al. Chimpanzee and human y chromosomes are remarkably divergent in structure and gene content. Nature 463, 536–539 (2010).

4. Hughes, J. F. et al. Strict evolutionary conservation followed rapid gene loss on human and rhesus y chromosomes. Nature 483, 82–87 (2012).

5. Soh, Y. Q. S. et al. Sequencing the mouse y chromosome reveals convergent gene acquisition and amplification on both sex chromosomes. Cell 159, 800–813 (2014).

6. Tomaszkiewicz, M. et al. A time- and cost-effective strategy to sequence mammalian Y chromosomes: An application to the de novo assembly of gorilla Y. Genome Res. 26, 530–540 (2016).

7. Simpson, J. T. et al. Detecting DNA cytosine methylation using nanopore sequencing. Nat. Methods 14, 407–410 (2017).

8. Charlesworth, B. & Charlesworth, D. The degeneration of Y chromosomes. Philos. Trans. R. Soc. B Biol. Sci. 355, 1563–1572 (2000).

9. Hughes, J. F. & Page, D. C. The Biology and Evolution of Mammalian Y Chromosomes. Annu. Rev. Genet. 49, 507–527 (2015).

10. Skaletsky, H. et al. The male-specific region of the human Y chromosome is a mosic of discrete sequence classes. Nature 423, 825–837 (2003).

11. Zhang, K. et al. Sequencing genomes from single cells by polymerase cloning. Nat. Biotechnol. 24, 680–686 (2006).

12. Berlin, K. et al. Assembling large genomes with single-molecule sequencing and locality-sensitive hashing. Nat. Biotechnol. 33, 623–630 (2015).

13. Chaisson, M. J. P. et al. Resolving the complexity of the human genome using single-molecule sequencing. Nature 517, 608–611 (2015).

14. Kuderna, L. F. K. et al. A 3-way hybrid approach to generate a new high-quality chimpanzee reference genome (Pan tro 3.0). Gigascience 6, 1–6 (2017).

15. Bickhart, D. M. et al. Single-molecule sequencing and chromatin conformation capture enable de novo reference assembly of the domestic goat genome. Nat. Genet. 49, 643–650 (2017).

16. Jain, M. et al. Nanopore sequencing and assembly of a human genome with ultra-long reads. Nat. Biotechnol. (2018). doi:10.1038/nbt.4060

17. Jain, M. et al. Linear assembly of a human centromere on the Y chromosome. Nat. Biotechnol. 36, (2018).

18. Auton, A. et al. A global reference for human genetic variation. Nature 526, 68–74 (2015).

19. Koren, S. et al. Canu: scalable and accurate long- - - read assembly via adaptive k - - - mer weighting and repeat separation. 1–35 (2016). doi:10.1101/gr.215087.116.Freely

20. Walker, B. J. et al. Pilon: An integrated tool for comprehensive microbial variant detection and genome assembly improvement. PLoS One 9, (2014).

21. Page, D. C., Harper, M. E., Love, J. & Botstein, D. Occurrence of a transposition from the X-chromosome long arm to the Y-chromosome short arm during human evolution. Nature 311, 119–123 (1984).

22. Alkan, C. et al. Personalized copy number and segmental duplication maps using next-generation sequencing. Nat. Genet. 41, 1061–1067 (2009).

23. Zook, J. M. et al. Extensive sequencing of seven human genomes to characterize benchmark reference materials. Sci. Data 3, 1–26 (2016).

24. Hughes, J. F. & Rozen, S. Genomics and Genetics of Human and Primate Y Chromosomes. Annu. Rev. Genomics Hum. Genet. 13, 83–108 (2012).

25. Nattestad, M. & Schatz, M. C. Assemblytics: A web analytics tool for the detection of variants from an assembly. Bioinformatics 32, 3021–3023 (2016).

26. Poznik, G. D. et al. Punctuated bursts in human male demography inferred from 1,244 worldwide Y-chromosome sequences. Nat. Genet. 48, 593–599 (2016).

27. Hernando, H. et al. The B cell transcription program mediates hypomethylation and overexpression of key genes in Epstein-Barr virus-associated proliferative conversion. 1–16 (2013).

28. Lukaszewski, A. J. et al. A chromosome-based draft sequence of the hexaploid bread wheat (Triticum aestivum) genome. Science (80-. ). 345, (2014).

29. Zimin, A. V. et al. The first near-complete assembly of the hexaploid bread wheat genome, Triticum aestivum. Gigascience 6, 1–7 (2017).

30. Neale, D. B. et al. Decoding the massive genome of loblolly pine using haploid DNA and novel assembly strategies. Genome Biol. 15, 1–13 (2014).

31. Nowoshilow, S. et al. The axolotl genome and the evolution of key tissue formation regulators. Nature 554, 50–55 (2018).

## Material and Methods References

32. Gribble, S. M., Ng, B. L., Prigmore, E., Fitzgerald, T. & Carter, N. P. Array painting: A protocol for the rapid analysis of aberrant chromosomes using DNA microarrays. Nat. Protoc. 4, 1722–1736 (2009).

33. Li, H. & Durbin, R. Fast and accurate short read alignment with Burrows-Wheeler transform. Bioinformatics 25, 1754–1760 (2009).

34. Kurtz, S. et al. Versatile and open software for comparing large genomes. Genome Biol. 5, R12 (2004).

35. Wu, T. D. & Watanabe, C. K. GMAP: A genomic mapping and alignment program for mRNA and EST sequences. Bioinformatics 21, 1859–1875 (2005).

36. Slater, G. S. C. & Birney, E. Automated generation of heuristics for biological sequence comparison. BMC Bioinformatics 6, 1–11 (2005).

37. Li, H. Genome analysis Minimap2: pairwise alignment for nucleotide sequences. (2018). doi:doi.org/10.1093/bioinformatics/bty191

